# Revisiting the use of dioxane as a reference compound for determination of the hydrodynamic radius of proteins by pulsed field gradient NMR spectroscopy

**DOI:** 10.1101/2023.06.02.543514

**Authors:** Emil E. Tranchant, Francesco Pesce, Nina L. Jacobsen, Catarina B. Fernandes, Birthe B. Kragelund, Kresten Lindorff-Larsen

**Affiliations:** Structural Biology and NMR Laboratory, The Linderstrøm-Lang Centre for Protein Science, and; REPIN, Department of Biology, University of Copenhagen, Copenhagen, Denmark

**Keywords:** IDP, ensemble, diffusion coefficient, 1,4-dioxane, hydrodynamic radius, radius of gyration, SAXS, lysozyme

## Abstract

Measuring the compaction of a protein or complex is key to understand the interactions within and between biomolecules. Experimentally, protein compaction is often probed either by estimating the radius of gyration (*R*_g_) obtained from small-angle X-ray scattering (SAXS) experiments or the hydrodynamic radius (*R_h_*) obtained for example by pulsed field gradient nuclear magnetic resonance (PFG NMR) spectroscopy. PFG NMR experiments generally report on the translational diffusion coefficient, which in turn can be used to estimate *R_h_* using an internal standard. Here, we examine the use of 1,4-dioxane as an internal NMR standard to account for sample viscosity and uncertainty about the gradient strength. Specifically, we revisit the basis for the commonly used reference value for the *R_h_* of dioxane (2.12 Å) that is used to convert measured diffusion coefficients into a hydrodynamic radius. We follow the same approach that was used to establish the current reference value for the *R_h_* by measuring SAXS and PFG NMR data for a set of seven different proteins and using these as standards. Our analysis shows that the current *R*_h_ reference value for 1,4-dioxane *R*_h_ (2.12 Å) is underestimated, and we instead suggest a new value of 2.27 Å ± 0.04 Å. Using this updated reference value results in a ∼7% increase in *R*_h_ values for proteins whose hydrodynamic radius have been measured by PFG NMR. We discuss the implications for ensemble descriptions of intrinsically disordered proteins and evaluation of effect resulting from for example ligand binding, posttranslational modifications, or changes to the environment.

## INTRODUCTION

Proteins are dynamic entities that exist in ensembles of states whose average properties vary depending on their sequence properties, context, and post-translational modifications. Folded proteins typically have a narrow distribution of conformations, whereas the structures of so-called intrinsically disordered proteins (IDPs) vary substantially across the ensemble (1). When characterizing the structure and interactions of proteins it is often advantageous to be able to probe the expansion of the protein or the size of the assembly it forms. Experimentally, this can for example be assessed via probing the radius of hydration, *R*_h_, and the radius of gyration, *R*_g_ (2, 3). *R*_h_ is often probed by pulsed field gradient nuclear magnetic resonance (PFG NMR) spectroscopy (4), fluorescence correlation spectroscopy, dynamic light scattering or, with a lower resolution, by size exclusion chromatography. *R*_g_ is most commonly probed by small-angle X-ray scattering (SAXS) (2, 5). As a robust and accurate determination of these two parameters is critical, and current estimates of *R*_h_ using PFG NMR rely on assumptions that for decades have mostly been left unexamined, we wanted to revisit one key step when determining the *R*_h_ of a protein by PFG NMR.

In PFG NMR, the location of the protein is “encoded” via a spatial field gradient and makes it possible to probe the translational diffusion coefficient (*D_t_*) and in turn estimate the *R*_h_. As PFG NMR reports on the conformationally averaged *D_t_*, and thus average *R*_h_, the technique is especially useful when studying IDPs and their interactions, where conformational ensemble compactness can provide structural information and interaction affinities (6, 7). In a standard PFG NMR one records a series of NMR spectra at varying gradient strengths, where the decay of the NMR peak intensities with increasing gradient strengths can be fitted to the Stejskal-Tanner equation (Eq. 1) (8), that relates *D_t_* to the measured peak intensities:

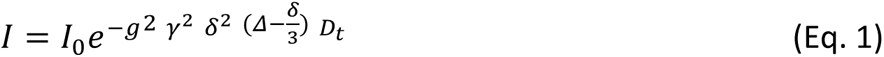

Here, *g* is the gradient strength, *γ* is the gyromagnetic ratio, *δ* is the length of the gradient, and *Δ* is the diffusion time. Assuming that the resulting translational diffusion of the nuclei is equal to that of the parent molecule, and the molecule diffuses as a spherical entity, the *R*_h_ of the selected peak can then be derived from the Stokes-Einstein relation (Eq. 2).

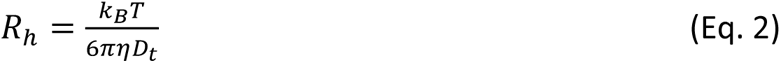

where *k*_B_ is the Boltzmann constant, *T* the temperature, and *η* the solvent viscosity; Eq. 2 can also be used as a definition of *R*_h_. Complications may, however, arise when determining the *R*_h_ this way. First, estimating *D_t_* on an absolute scale requires careful calibration of the field gradient.

Second, the solvent viscosity is sensitive to type of solvent, solute and sample conditions. Thus, the buffer composition, temperature, and protein concentration may all affect viscosity and should be carefully controlled. Furthermore, any added D_2_O used to lock the NMR frequency must be corrected for as there is a slight difference in viscosity between H_2_O and D_2_O (7). As the solvent viscosity is difficult to measure and control precisely, deriving reliable *R*_h_ of a protein from a selected peak just from its translational diffusion alone is difficult and may be imprecise. Instead, one often uses an internal reference compound that is added to the NMR sample. By knowing the *R*_h_ of the reference compound (9), the ratio of the *D_t_* between the reference compound and the protein can be used to estimate the *R*_h_ of the protein according to Eq. 3.

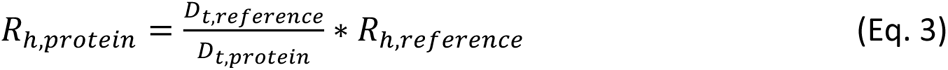

Using an internal reference removes the need for accurate calibration of the gradient and measurement of the viscosity. Often, the reference compound of choice is 1,4-dioxane. 1,4– dioxane provides a single, easily discernible, _1_H NMR peak at approx. 3.75 ppm and has been found not to interact substantially with several proteins at experimentally used concentrations (10). While other reference compounds such as *α-* or *β-*cyclodextrin are sometimes suggested in place of 1,4-dioxane (11), cyclodextrins have also been shown to interact with proteins (12, 13), just as they contribute with many more signals in the NMR spectrum overlapping with those of the proteins.

Using 1,4-dioxane as a viscosity reference requires that its *R*_h_ is known and that it is insensitive to environmental changes. Early use of 1,4-dioxane as a reference in PFG NMR established the *R*_h_ of the molecule to be 2.12 Å, and this value has since been used as a reference when using PFG NMR to determine the *R*_h_ of proteins (14). The reference value for dioxane was determined as described above, but instead using a protein molecule as reference. Specifically, Wilkins et al performed PFG NMR experiments on a solution of 1,4-dioxane and hen egg white lysozyme (HEWL), where instead of an unknown *R*_h_ of the protein in Eq. 3, the *R*_h_ of dioxane was unknown and the *R*_h_ of the HEWL set to 19.8 Å. This *R*_h_ value originated from an earlier study where batch SAXS experiments on HEWL provided an experimental *R*_g_ for natively folded HEWL of 15.3 ± 0.2 Å (15). By assuming the ratio between *R*_g_ and *R*_h_, *ρ*, for a globular protein such as HEWL to be that of a solid sphere (16), the *R*_h_ of HEWL was obtained with the *R*_g_ from SAXS using equation 4:

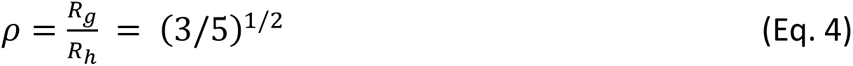

Using this approach and experimental batch SAXS data of HEWL, the authors (14) estimated the *R*_h_ of natively folded HEWL to be 19.8 Å. The ratio of the translational diffusion coefficients between HEWL and 1,4-dioxane was estimated from the PFG NMR data to be 9.33 (17), leading to an *R*_h_ of 1,4-dioxane of 2.12 Å.

Recently, a community study addressed the reproducibility of SAXS experiments between proteins and between instruments (18). Generally, for the studied set of proteins, experimental reproducibility and subsequent consensus curves were observed. However, batch SAXS experiments on HEWL specifically showed significant variability between experiments. Considering that the original derivation of the *R*_h_ of 1,4-dioxane was directly derived based on a batch SAXS measurement of HEWL, any uncertainty in the *R*_g_ of HEWL would result in uncertainty in the 1,4– dioxane *R*_h_ of 2.12 Å and hence impact *R*_h_ measurements of proteins using PFG NMR and dioxane referencing.

With this possible uncertainty on the *R*_h_ of 1,4-dioxane in mind, we decided to revisit the foundation of this reference value by exploring a larger set of proteins, using the same approach as originally done in (14). We estimate the average and uncertainty of the *R*_h_ of 1,4-dioxane using a set of seven folded proteins for which we measured both translational diffusion coefficients by PFG NMR and determined the *R_g_* by batch SAXS measurements. By considering the quality of the recorded data, we find that the established *R*_h_ value of 1,4-dioxane is slightly underestimated. Our data suggests that it should be increased by approximately 7% compared to the previous value, and we propose an updated standard *R*_h_ value of 2.27 ± 0.04 Å of 1,4-dioxane, which would also result in a 7% increase in derived protein *R*_h_ using Eq. 4.

## MATERIALS AND METHODS

### Protein purifications

Protein samples used in this work were either prepared from bought lyophilized powder stocks or from frozen, pre-purified stocks. Proteins from purchased powder-stocks were equine myoglobin (Sigma-Aldrich), bovine ribonuclease A (RNaseA) (GE Healthcare), and HEWL (Sigma-Aldrich). Prepurified proteins include S100A13, prolactin, ACBP_Y73F_, and 14-3-3 ζ. Purification of prolactin was performed as described in (19). Purification of ACBP_Y73F_ was performed as described in (20).

For S100A13, *E. coli* cells (BL21 DE3) (Biolabs) were transformed with pET-24a plasmid coding for His_6_-SUMO S100A13 (UniProt Q99584) and grown in high salt LB-broth medium (Sigma Aldrich). Cells were grown until an OD_600_ of 0.6-0.8 and expression induced with 0.1 mM isopropyl-β-D-1-thiogalactopyranoside (IPTG). After 4h growth at 37 ^°^C, cells were harvested by centrifugation at 5000 x g for 15 min and stored at –20 ^°^C. Cells were lysed in 50 mM Tris pH 8.0, 150 mM NaCl, 2 mM CaCl_2_ through a French press cell disrupter at 25,000 psi (Constant Systems Ltd.) followed by centrifugation at 20,000 x g where the clear lysate was subsequently loaded onto a 5 mL Ni-NTA Sepharose column (GE Healthcare) equilibrated with 50 mM Tris, pH 8.0, 150 mM NaCl, 2 mM CaCl_2_. His_6_-SUMO S100A13 was eluted with 50 mM Tris pH 8.0, 150 mM NaCl, 500 mM Imidazole followed by overnight dialysis where 0.1 mg His-tagged ULP1 and 1 mM DTT was added to cleave the SUMO-tag. The sample was purified further using a reverse Ni-NTA step removing non-cleaved protein, ULP1 and SUMO followed by removal of DNA on a 1 mL Heparin column with 50 mM Tris, pH 7.4, 2 mM CaCl_2_ and 50 mM Tris pH 7.4, 1 M NaCl, 2 mM CaCl_2_ For a final step, the sample was run on a Superdex 75 10/300 (GE Healthcare) in 50 mM Tris pH 7.4, 150 mM NaCl, fractions concentrated and stored at –20 ^°^C.

The 14-3-3ζ protein was expressed from a modified pET-24a vector designed to encode an N-terminal His_6_-SUMO tag, which was to be cleaved using ubiquitin-like protein protease 1 (ULP1). The plasmid was transformed into NiCo21(DE3) competent *E. coli* cells (New England BioLabs) grown in LB medium containing 50 µg/mL kanamycin and the fusion protein expression was induced with 0.5 mM IPTG for 4-5 hours before harvesting cells by centrifugation at 5000xg, 15 min, 4 °C. The pellet was lysed in lysis/equilibration buffer (20 mM Bis-Tris pH 7.2, 10 mM Imidazole, 150 mM NaCl, 5 mM β-mercaptoethanol (bME)) using a French pressure cell disrupter (25 kpsi; Constant Systems Ltd, Daventry, UK), and the lysate was cleared by centrifugation at 20,000xg for 45 min at 4 °C. The His_6_-SUMO-14-3-3ζ fusion protein was purified by immobilized metal affinity chromatography (IMAC) using Ni Sepharose 6 Fast Flow resin (5 mL; GE Healthcare) with standard IMAC procedures of sample application, high salt (1 M NaCl) washing step and imidazole elution. The eluted sample was dialyzed towards 2 L of buffer A (20 mM Bis-Tris pH 6.5, 5 mM bME) before the sample was applied to a 1 mL HiTrap Heparin HP column (Cytiva). The column was washed with 15 mL of buffer A before the fusion protein was eluted with a linear two-step gradient of 0-30% over 3 mL and 30-100% over 20 mL of buffer B (20 mM Bis-Tris pH 6.5, 1 M NaCl, 5 mM bME). The His_6_-SUMO tag was cleaved off by supplementing the sample with 0.1 mg ULP1 and 2 mM DTT for at least 3 hours. This sample was re-applied to the IMAC column to remove the His_6_-SUMO-tag and ULP1, and the flow through containing pure 14-3-3ζ was collected.

### NMR and SAXS sample preparations

To prepare samples from purified proteins or lyophilized protein stocks, each protein was applied to a size exclusion chromatography (SEC) column (Superdex 75 10/300 GL, *Cytiva*) in a 20 mM sodium phosphate buffer, pH 7.4, 150 mM NaCl mounted on an Äkta Purifier system. Each collected sample was then analysed by SDS-PAGE to verify the purity of the protein. The samples for PFG NMR experiment were prepared with the following protein and concentrations: HEWL 200 µM, RnaseA 300 µM, myoglobin 400 µM, S100A13 150 µM, ACBP_Y73F_ 200 µM, prolactin 300 µM, and 14-3-3 150 µM. DSS was added to a final concentration of 25 µM, D_2_O to a final concentration of 10% (v/v), and 1,4-dioxane to a final concentration between 0.04-0.06% (v/v). Sample volumes were either 100 µL, 350 µL, or 500 µL, depending on the use of 3 mm Shigemi, 5 mm Shigemi, or 5 mm glass single-use NMR tubes (Bruker), respectively. Samples for SAXS experiments were prepared by concentrating the proteins after SEC to multiple samples of 1, 2, and 3 mg/ml in the 20 mM sodium phosphate buffer also used in the SEC. A pure buffer solution was also prepared for buffer subtraction of the background scattering.

### Pulsed field gradient NMR Spectroscopy

PFG NMR experiments were performed on a Bruker Avance III HD 600 MHz spectrometer equipped with a Bruker proton-optimized quadruple resonance NMR ‘inverse’ QCI cryoprobe. Each PFG NMR experiment was preceded by a 1D ^1^H-spectrum used for referencing the spectra to the DSS peak at 0 ppm. Translational diffusion coefficients of proteins were determined by fitting peak intensity decays in the methyl and methylene region (2.5-0.5 ppm) to the Stejskal-Tanner equation (8, 9). The 1,4-dioxane translational diffusion coefficients were fitted to the intensity decay of the 1,4-dioxane peak at 3.75 ppm. Integration of selected peaks was performed by Bruker Dynamics Center v2.5.6. For every PFG NMR experiment, 64 scans were recorded with a gradient strength interval from 2% to 98% (*γ*=26752 rad s^-1^ Gauss^-1^) with a diffusion time (Δ) of 200 ms and a gradient length (δ) of 2 ms. Baseline correction and assessment of the Stejskal Tanner fitting intervals were performed in Bruker Topspin 3.6.2 and Dynamics Center v2.5.6, while final fitting of the translation diffusion was performed in GraphPad Prism 8.2.1.

### SAXS experiments

SAXS experiments were performed at the CPHSAXS Facility, University of Copenhagen, on a Xenocs© BioXolver L with a wavelength of λ = 1.34 Å. Primary data reduction was made in BIOXTAS RAW. For each sample protein the optimal protein concentration yielding a high signal to noise ratio while avoiding aggregation was chosen by inspecting the low-*q* region of the scattering curves by Guinier analysis. Each scattering curve consisted of an accumulation of 10 measurements on each sample. Subsequent data analysis was performed in the ATSAS 3.0.5 Primus suite (21), with the “merge” function used on multiple scattering curves for each sample protein. Merged scattering curves of *n* SAXS experiments for each protein sample were then used for Guinier derivation of the protein *R*_g_. The Primus “AutoRg” function was used to determine *q*-range for *R*_g_ derivation. Pair distance distribution plots of the consensus curves were also calculated in the ATSAS 3.0.5 Primus suite with the *D*_max_ value being set based on a qualitative assessment of the P®-function reaching 0.

### Radius of hydration and gyration calculation from atomic coordinates

The anhydrous R_g_ from the atomic coordinates of proteins was calculated as the mass-weighted average distance of each atom from the protein’s centre of mass. For a better comparison to R_g_ values obtained from experimental SAXS data, we also used the WAXSiS webserver (22, 23) to explicitly include a water layer around the protein, calculate the SAXS profile of the envelope and fit an R_g_ from this with the Guinier approach. The *R*_h_ was calculated with the HullRadSAS software (24). For PDB entries that are NMR ensembles, the *R*_g_ and *R*_h_ were calculated on all conformers of the ensembles and then averaged as 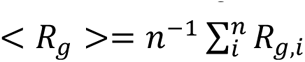 and 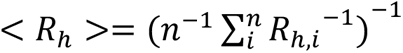. Missing C-terminal residues in the 14-3-3 ζ structure were added with MODELLER (25, 26) prior to calculating *R*_h_ and *R*_g_.

### Rh determination of 1,4-dioxane

To determine the *R*_h_ of 1,4-dioxane, the ratio of *D_t_* between the sample protein and 1,4-dioxane was used in Eq. 3 alongside the assumed protein *R*_h_ values calculated from experimental SAXS *R*_g_ values and Eq. 4. To ensure an accurate estimate of the final *R*_h_ value of 1,4-dioxane, the data quality was factored in by weighing the data by its quality using a χ^2^-approach. The χ^2^-value was calculated by first calculating the observed ratios of diffusion coefficients from PFG NMR, and then by subtracting the expected ratios of the diffusion coefficients from SAXS *R*_g_ derived *R*_h_ values and an assumed 1,4-dioxane *R*_h_ value spanning an interval of 2.0 to 2.5 Å. The deviation in the diffusion ratio from the observed NMR-data and SAXS-derived expected data was then divided by the sum of the squared experimental standard error of fits from both PFG NMR and SAXS, leaving a χ^2^-value representing the data quality and fit to different possible *R*_h_ values for 1,4-dioxane spanning 2.0 to 2.5 Å. We thus calculated χ^2^ using the following equation for different estimates of the *R*_h_ value of dioxane.

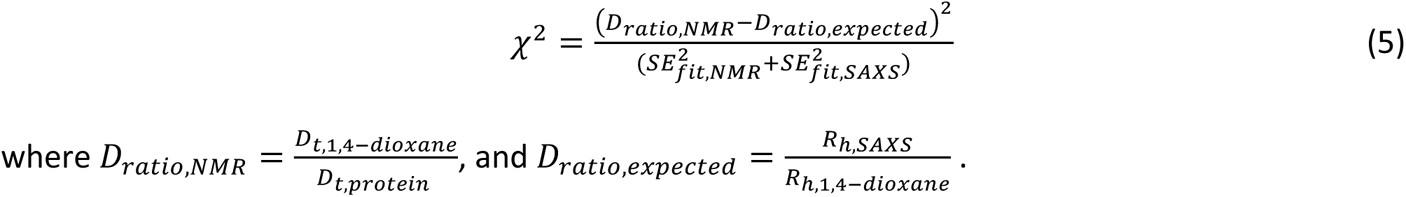

### Data availability

Experimental data, processed data, and scripts to reproduce the content of this work are available at: https://github.com/KULL-Centre/_2023_dioxane-tranchant/.

## RESULTS & DISCUSSION

### Proteins and experimental measurements

To establish a foundation for determining the *R*_h_ of 1,4-dioxane we chose a subset of seven globular proteins varying in size, and recorded PFG NMR and batch SAXS experiments for each protein at different concentrations. The seven proteins were chosen based on availability, ease of experimental work (e.g., solubility and stability), size and roughly spherical shapes based on earlier experimental structures, as to adhere to the assumption made in Eq. 4. The seven proteins were HEWL, RNaseA, myoglobin, S100A13 (dimer), acyl-CoA-binding protein Y73F (ACBP_Y73F_), prolactin and the 14-3-3 ζ dimer (see Fig. 1 & Table 1). All proteins were checked for purity and homogeneity by an SDS-PAGE and size-exclusion chromatography prior to NMR and SAXS data acquisition (see supplemental Fig. 1).

**Fig. 1:**
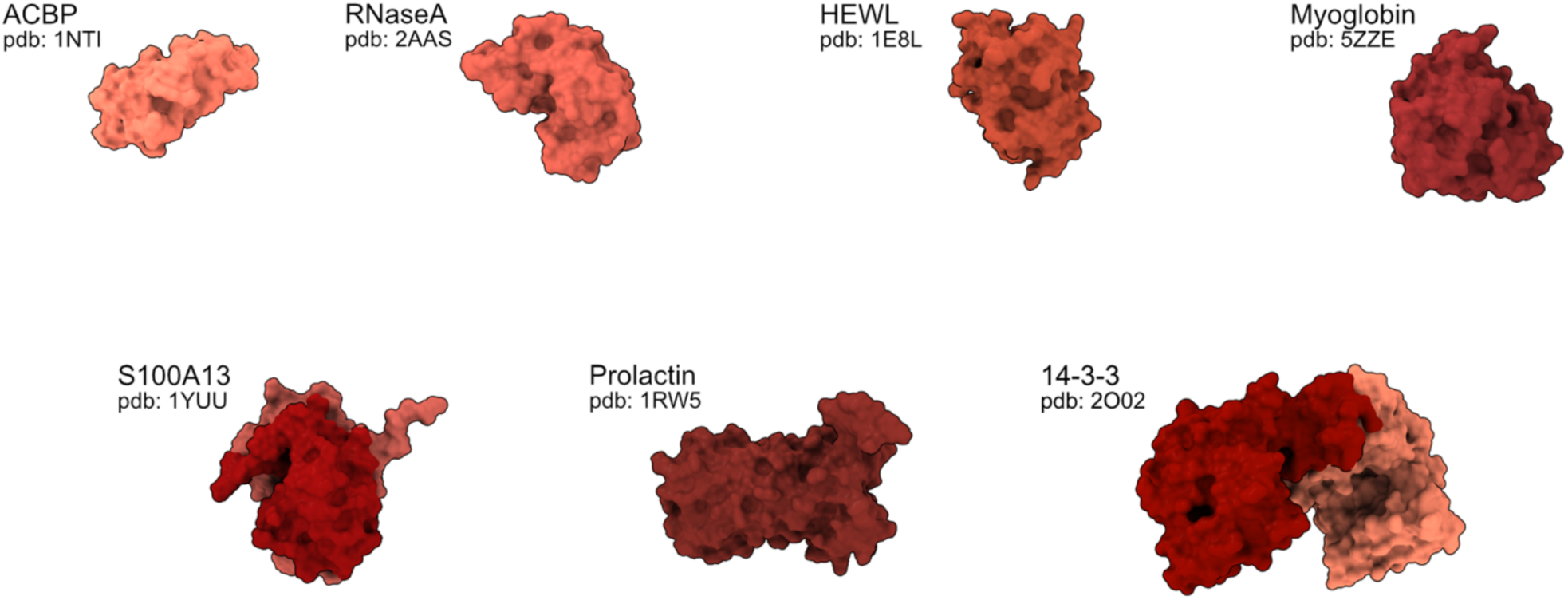
Surface contour representations of the seven sample proteins. Model 1 of the ensemble in the respective PDB entries are shown. Proteins are arranged in order of molecular weight and shown on comparable scales. The structures used are from PDB ID 1NTI (20), 2AAS (27), 1E8L (28), 5ZZE (29), 1YUU (30), 1RW5 (31), 2O02 (32).

**Table 1.** Overview of protein properties.

| PROTEIN | ACBP<sub>Y73F</sub> | RNASEA | HEWL | MYOGLOBIN | S100A13 | PROLACTIN | 14-3-3Z |
| --- | --- | --- | --- | --- | --- | --- | --- |
| ORGANISM | *B. taurus* | *B. taurus* | *G. gallus* | *E. caballus* | *H. sapiens* | *H. sapiens* | *H. sapiens* |
| SOURCE | *E. coli* expression | GE Healthcare | Sigma Aldrich | Sigma Aldrich | *E. coli* expression | *E. coli* expression | *E. coli* expression |
| UNIPROT ID (SEQUENCE) | P07108 (2–87, Y73F) | P61823 (27–150) | P00698 (19–147) | P68082 (2–154) | Q99584 (1–98) | P01236 (29–227) | P63104 (1–245) |
| PDB | 1NTI | 2AAS | 1E8L | 5ZZE | 1YUU | 1RW5 | 2O02* |
| MW | 9.9 kDa | 13.7 kDa | 14.3 kDa | 17.1 kDa | 22.9 kDa dimer | 22.9 kDa | 54 kDa dimer |

\*lacks 15 C-terminal residues compared to the WT used in these experiments.

We first probed the diffusion coefficients of each protein using PFG NMR experiments with the addition of 1,4-dioxane as an internal standard. After picking peaks corresponding to either 1,4-dioxane or protein, we fitted the intensity decays as a function of the gradient strength to the Stejskal Tanner equation (Eq. 1) and the *D*_t_ of protein and dioxane was estimated (Fig. 2 and Table 2).

**Fig. 2:**
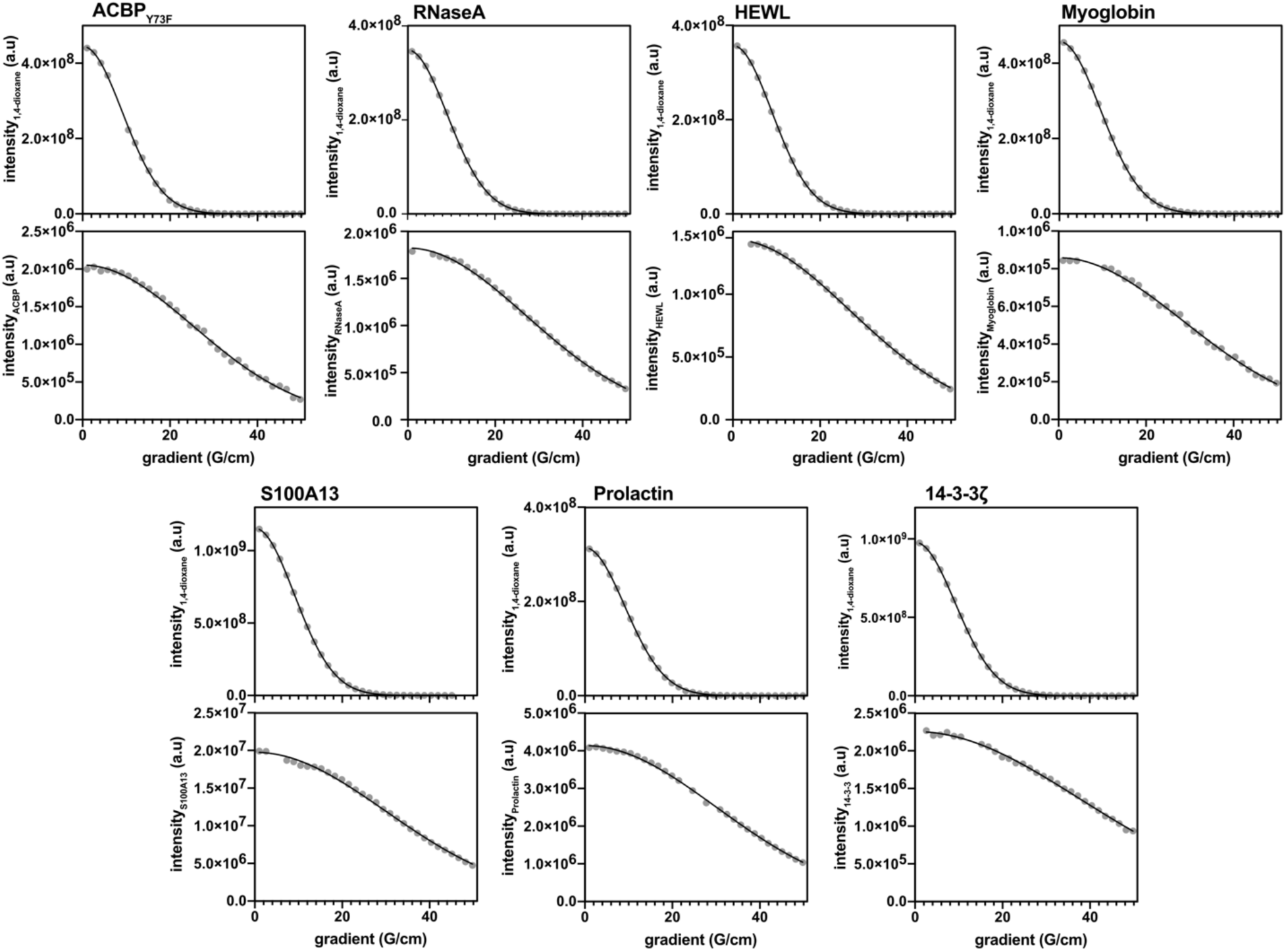
Peak intensity decays as a function of gradient strength in PFG NMR experiments. Intensity decays for seven samples containing both 1,4-dioxane (top) and protein (bottom). Samples are shown in the following order: ACBPY73F, RNaseA, HEWL, myoglobin, S100A13, prolactin, 14-3-3ζ. Differences in peak intensity of 1,4-dioxane between samples are due to variations in concentration and differences in automatic selection of integration intervals performed in Bruker Dynamics Center.

**Table 2.**
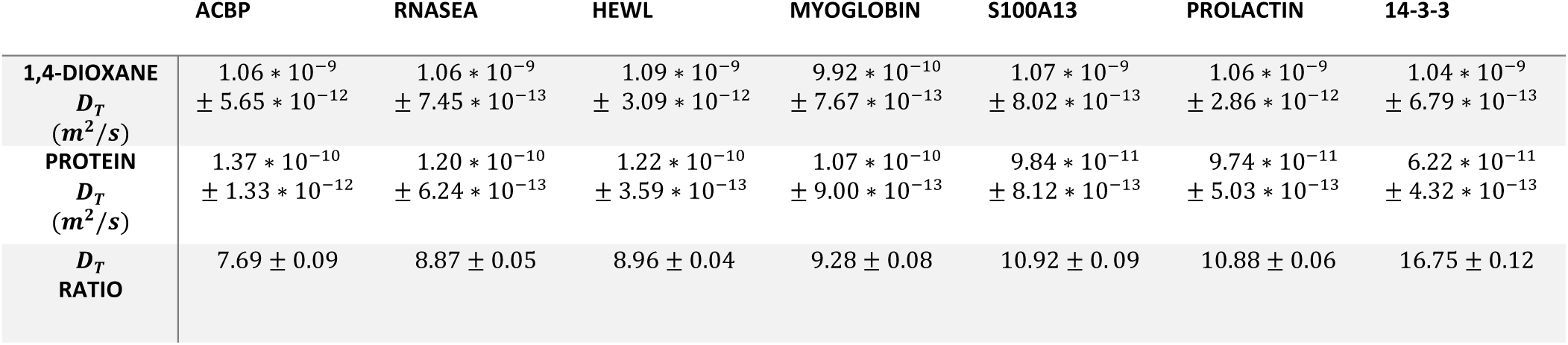
Diffusion coefficients of 1,4-dioxane and seven proteins and their ratio.

While the 1,4-dioxane data fitted nicely to the Stejskal Tanner equation, the protein data showed greater variation. In general, the ratio of the diffusion coefficients between protein and 1,4-dioxane increased as the molecular weight of the protein increased, as would be expected. Differences in total intensity of 1,4-dioxane was observed, which was expected as the 1,4-dioxane concentration in some of the samples was slightly different (0.06% in S100A13 and 14-3-3, 0.04% in ACBP, HEWL, RNaseA, prolactin, and myoglobin). However, the observed difference in 1,4-dioxane intensity was also observed in experiments on triplicates of identical samples (supplemental figure 2), likely attributed to the automatic selection of integration area by the Dynamics Center software when peak picking.

Notably, the ratio of the diffusion coefficients between HEWL and 1,4-dioxane in our experiments was lower (8.96 ± 0.05) than in the original work where the *R_h_* of 1,4-dioxane was estimated (9.33) (14, 17). This difference could be explained either by difference in pH or protein concentration between measurements, as the diffusion coefficient ratio of 9.33 was measured at pH 2, and with a HEWL concentration of ca. 1 mM, whereas our experiments were recorded at neutral pH and with 200 µM of HEWL. To test whether the observed variation in diffusion ratio between our experiments and the original experiments could be explained by instrumental error, we recorded technical triplicate measurements of HEWL and RNaseA samples prepared in the same way from the same protein stock (supplemental Fig. 2 and Table 2). These results showed an approximately 1.4% variance in the ratio of the diffusion coefficients between technical replicas, smaller than the difference between 8.96 and 9.33.

Next, we recorded multiple SAXS curves for each protein at optimal protein concentrations and used the “merge” function in the ATSAS suite to derive a protein consensus curve (Fig. 3). We then performed a Guinier analysis of each consensus curve to estimate *R*_g_ for each protein to help minimize effects from aggregation on the scattering curves that could otherwise increase variance between measurements. The quality of the consensus scattering curves was evaluated by examining the pair distance distribution plots of each curve (supplemental Fig. 3).

**Fig. 3:**
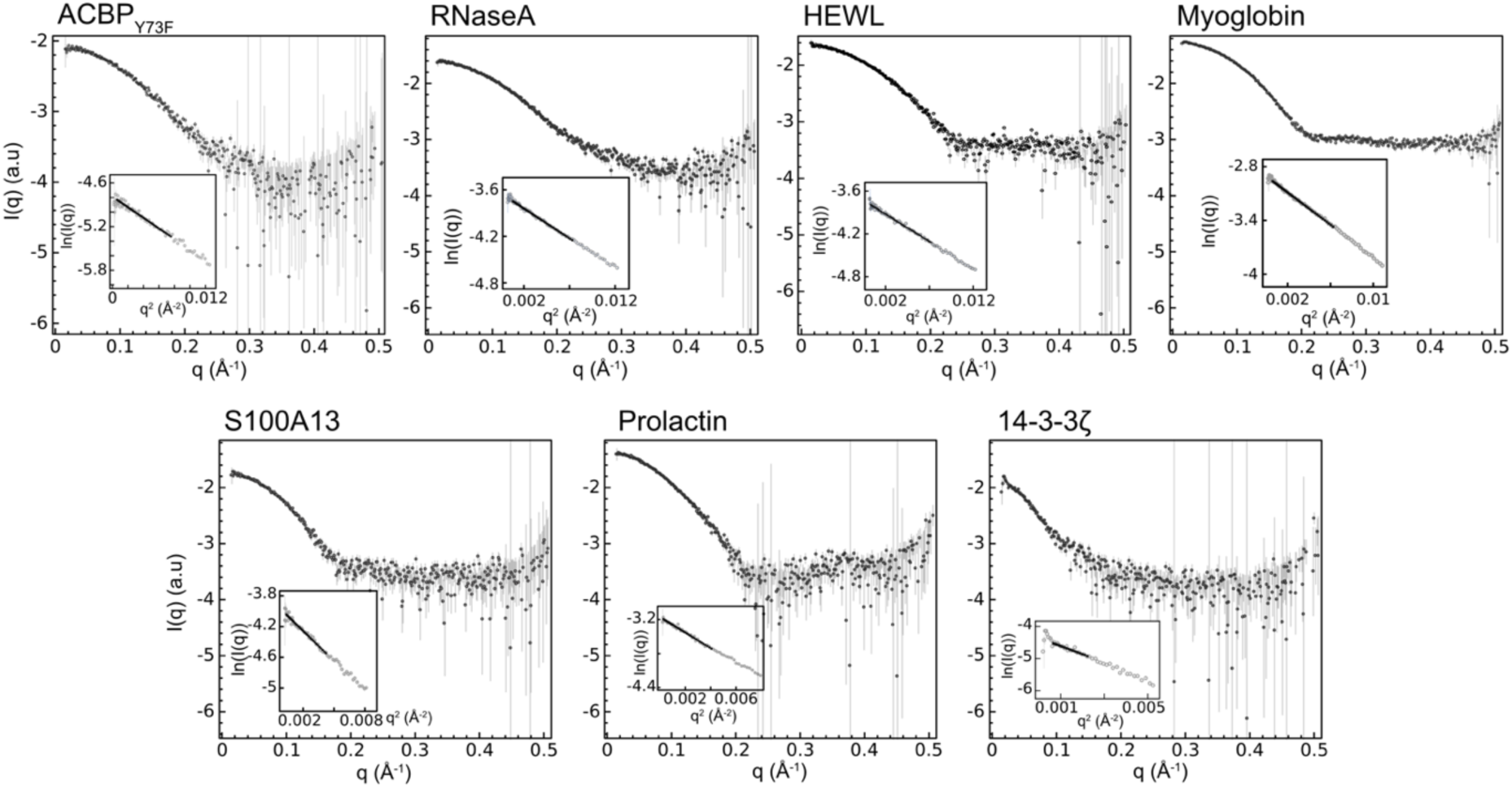
Consensus scattering curves of experimental scattering curves of each of the seven proteins. Total scattering curves used in each consensus curve: ACBP (from 2 measurements), RNaseA (4 measurements), HEWL (5 measurements), myoglobin (6 measurements), S100A13 (5 measurements), prolactin (3 measurements), and 14-3-3 (3 measurements). For each panel, the insert shows the low-*q* region on a logarithmic scale and the linear fit used for the Guinier analysis.

We estimated the *R*_g_ values of the proteins by Guinier analysis and from the pair distance distribution plots (supplemental table 3); only values from Guinier analysis were used for estimating the 1,4-dioxane *R*_h_. As expected, protein *R*_g_ increased as the molecular weight increased. Comparing the HEWL Guinier derived *R*_g_ of 15.16 ± 0.08 Å to the *R*_g_ used for the original 1,4-dioxane *R*_h_ derivation of 15.3 ± 0.2 Å (15), or to the mean batch-SAXS Rg of HEWL of 15.32 ± 0.81 Å from (18), measured in the presence of 2 M urea, revealed no substantial differences in the mean value.

We then used the measured *R*_g_-values for the seven proteins and the *D*_t_ ratios from PFG NMR to the *R*_h_ for 1,4-dioxane using Eq. 4 by estimating the protein *R*_h_-values from Eq. 3. This approach yielded an estimated 1,4-dioxane *R*_h_ from each protein data set (Table 4).

**Table 4:** Experimentally derived diffusion coefficient ratios from PFG NMR, experimentally derived *Rg* values by Guinier analysis from batch SAXS, estimated protein *R*h values from the SAXS data and resulting estimated *R*h values from 1,4-dioxane. All reported errors are propagated standard errors of the original fits.

|  | ACBP | RNASEA | HEWL | MYOGLOBIN | S100A13 | PROLACTIN | 14-3-3 |
| --- | --- | --- | --- | --- | --- | --- | --- |
| $D_T$<br>RATIO | $7.69 \pm 0.09$ | $8.87 \pm 0.05$ | $8.96 \pm 0.04$ | $9.28 \pm 0.08$ | $10.92 \pm 0.09$ | $10.88 \pm 0.06$ | $16.75 \pm 0.12$ |
| PROTEIN<br>$R_g$ (Å)<br>(Guinier) | $14.79 \pm 0.18$ | $15.20 \pm 0.08$ | $15.16 \pm 0.08$ | $16.41 \pm 0.04$ | $19.77 \pm 0.24$ | $20.29 \pm 0.16$ | $27.47 \pm 1.00$ |
| ESTIMATED<br>PROTEIN<br>$R_h$ (Å) | $19.09 \pm 0.23$ | $19.62 \pm 0.10$ | $19.57 \pm 0.10$ | $21.19 \pm 0.05$ | $25.52 \pm 0.31$ | $26.19 \pm 0.21$ | $35.46 \pm 1.29$ |
| DERIVED<br>1,4-DIOXANE<br>$R_h$ (Å) | $2.48 \pm 0.06$ | $2.21 \pm 0.02$ | $2.18 \pm 0.02$ | $2.28 \pm 0.02$ | $2.34 \pm 0.05$ | $2.41 \pm 0.03$ | $2.12 \pm 0.09$ |

Depending on the protein used as a standard, we arrived at different estimates of the *R*_h_ values for 1,4-dioxane ranging between 2.12 Å to 2.48 Å. The original estimate of *R*_h_ = 2.12 Å lies at the edge of this interval of *R*_h_ values but appears to be an underestimate as six of the seven protein data sets gave larger values. Taking the average 1,4-dioxane *R*_h_ from these data, and factoring in the errors for each protein set, we arrive at a weighted average of 2.27 Å, with an error estimated by bootstrapping to be 0.02 Å.

### Examining the relationship between R_g_ and Rh for proteins

It is possible that differences in the *R*_h_ values estimated for 1,4-dioxane from the SAXS and NMR data across the seven proteins can in part be explained by deviation of the protein shapes from sphere of uniform mass distribution. As a consequence, the relationship between *R*_g_ and *R*_h_ may not be equal to ρ=(3/5)^1/2^, as assumed in Eq. 4. To examine whether this assumption is reasonable for the seven proteins, we used both the crystallographic and solution structures of each of the seven proteins to calculate *R*_g_ and *R*_h_, and compared their ratio to the assumed value of ρ=(3/5)^1/2^ (Fig. 4) (23). Deviations from an ideal value of ρ would affect our analysis and the estimates of the *R*_h_ value for 1,4-dioxane presented above. We selected the seven proteins to have roughly spherical shapes and we expect small deviations from ρ=(3/5)^1/2^ to average out over the dataset. Overall, we observe the calculated *R*_g_/*R*_h_ values to be close to ρ=(3/5)^1/2^, with a small underestimation on average. As the *R*_g_ in Fig. 4 was calculated only for the protein coordinates, we subsequently included the contribution of the hydration layer to the calculated *R*_g_ to investigate if the apparent underestimation of *R*_g_/*R*_h_ could be accounted for by considering the hydration of the proteins (supplemental Fig. 4) (24). This, however, led to a large overestimation of *R*_g_/*R*_h_ compared to ρ=(3/5)^1/2^. We examined whether the deviation from to ρ=(3/5)^1/2^ could be explained by deviations from a spherical shape by calculating the aspherity of the proteins (33); however, we find no strong correlation between aspherity and observed deviation (supplemental Fig. 5).

**Fig 4:**
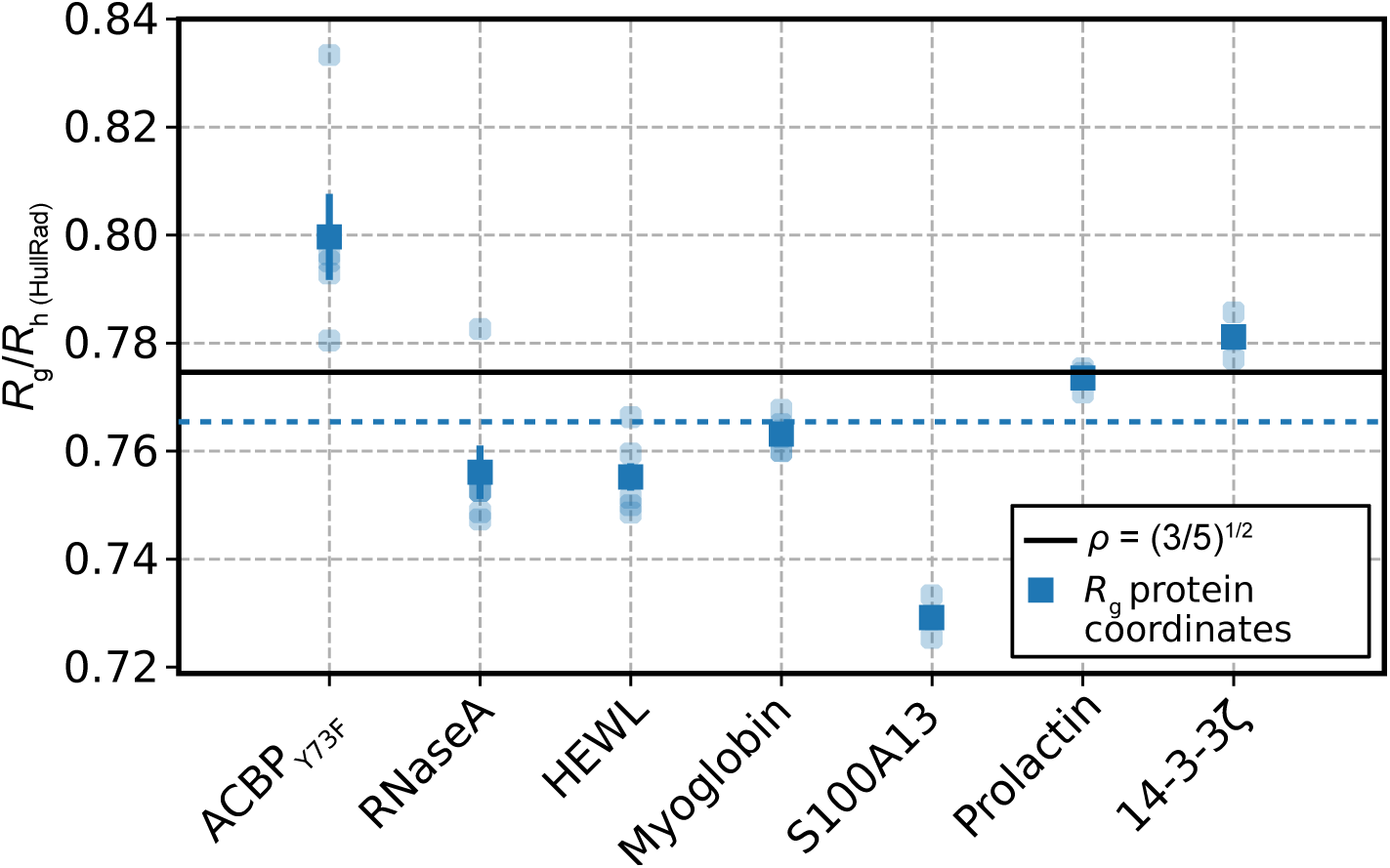
Ratios of *R*g/*R*h obtained from 3D structures. For each protein, three to five 3D structures were used, with the following PDB codes for each protein: HEWL (1e8l (28), 1dpx (34), 6abn (35), 5a3e (36)), RNaseA (2aas (27), 1fs3 (37), 1jvt (38), 4ooh(39), 1kf7 (40)), myoglobin (5zze (29), 1wla (41), 4dc8 (42), 5d5r (43), 5cn4 (43)), S100A13 (1yuu (30), 2h2k (44), 2egd (45)), ACBP (1nti (20), 1hb6 (46), 1hb8 (46), 2abd (47)), prolactin (1rw5 (31), 2q98 (48)), 14-3-3ζ (2o02 (32), 1qja (49), 1gjb (50)). AlphaFold structures (51, 52) were also included for each protein. Individual ratios are shown as partially transparent markers. For each protein we calculate the mean ration over different structures and the standard error of the mean (solid markers). Dashed blue line represents the calculated average value of the *R*g/*R*h.

### Factoring in the data quality in calculating the expected R*_h_* of 1,4-dioxane

As observed in the difference of the 1,4-dioxane *R*_h_ calculated from the different protein data sets, the choice of protein can affect the outcome. We therefore performed a global analysis of all seven protein data sets to help minimize protein-specific effects on the estimated value of *R*_h_ for 1,4-dioxane. For this, we calculated a χ^2^ value (Eq. 5) between the measured ratio of diffusion coefficients and the value expected depending on (i) the estimated *R*_h_ for each protein and (ii) the *R*_h_ for 1,4-dioxane (Eq. 3). With this approach, we can assess an interval of likely 1,4-dioxane *R*_h_ values while taking into account the errors on the measured *D*_t_ ratios and the estimated value for the *R*_h_ for the seven proteins (from SAXS and ρ=(3/5)^1/2^) (16).

The resulting plot of χ^2^ vs. the *R*_h_ for 1,4-dioxane (Fig. 5) shows a minimum around *R*_h_ of 2.27 Å and, as discussed above, suggests that the previously determined value of 2.12 Å is an underestimate (dashed line in Fig. 5). To estimate an error on the χ^2^-estimated 1,4-dioxane *R*_h_ of 2.27 Å, we performed a leave-one-out analysis which provided an error estimate of 0.02 Å, corresponding with the earlier weighted average 1,4-dioxane *R*_h_ and bootstrap error estimation. We also estimated an error using the technical replicate measurements of HEWL and RNaseA in both PFG NMR and SAXS (supplemental table 1 and table 2, respectively). From the technical replicates, we found a 1.4% and 1.1% error across samples measured by PFG NMR and SAXS Guinier analysis, respectively. Propagating these relative errors from the technical replicates to the χ^2^-estimated 1,4-dioxane *R*_h_, we determine the *R*_h_ of 1,4-dioxane to be 2.27 ± 0.04 Å.

**Fig. 5:**
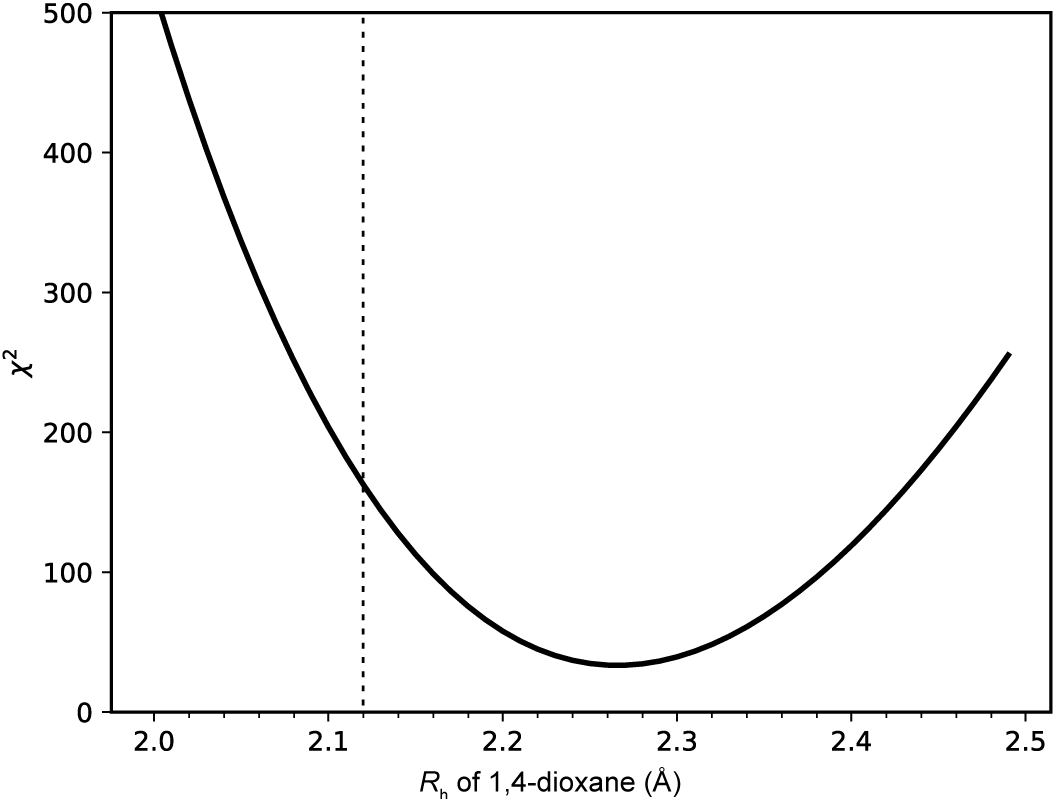
χ^2^-value for the seven protein data sets calculated as described in Eq. 5 for different possible *R*h values for 1,4-dioxane. The dashed vertical line highlights the *R*h value of 1,4-dioxane of 2.12 Å determined in (14).

### Impact of an increase in the size of the hydrodynamic ratio of 1,4-dioxane

With an added uncertainty to the estimated 1,4-dioxane *R*_h_, the above results suggest a ∼7% (2.27 Å/2.12 Å=1.07) increase in *R*_h_ compared to the commonly used reference. This change in reference value can be accounted for when examining previously published PFG NMR data that have used 1,4-dioxane as a reference by increasing the derived protein *R*_h_ by 7% as well. For example, we previously reported the *R*_h_ of prothymosin-α to be 28.9 ± 0.8 Å using PFG NMR and with 1,4-dioxane as a reference with the *R*_h_ set as 2.12 Å (53). Using the updated reference value for dioxane and propagating the errors, the re-estimated *R*_h_ would be 31 ± 1 Å. Often, *R*_h_ values from PFG NMR are used to track changes in protein dimensions following a selected perturbation. In these cases, the increase in 1,4-dioxane *R*_h_ would not be an issue, as the relative changes are unaffected by the increase in the absolute *R*_h_. An example of this would be PFG NMR experiments studying a protein oligomeric state at different concentrations, by deriving the oligomeric state from the observed *R*_h_ at different concentrations (54).

### Consequences for estimating the conformational ensembles of IDPs

One important consequence of the results presented here is that they affect our understanding of how to compare conformational ensembles with experimental measurements of *R*_h_. We recently compiled a list of the *R*_h_ of eleven IDPs as measured by PFG NMR, and used these together with SAXS experiments to evaluate different models to calculate *R*_h_ from conformational ensembles of IDPs (55). Our work led to the conclusion that, among the different approaches tested, the Kirkwood-Riseman equation (56) resulted in the best agreement between computational models and the *R*_h_ values measured by PFG NMR. We noted, however, how possible inaccuracies in the reference *R*_h_ of 1,4-dioxane would affect our results, by changing the experimentally determined *R*_h_ values and leading to a different conclusion.

As we here have shown that the *R*_h_ of 1,4-dioxane was previously underestimated, we re-examine the conclusion of our previous work considering the new reference value for the *R*_h_ of 1,4-dioxane. Therefore, we first increased the experimentally derived *R*_h_ values for the eleven IDPs by 7% (table S4) and applied an uncertainty of 2.1% (average relative error associated with the *R*_h_ of the IDPs) to the corrected *R*_h_ values, as we previously proposed to provide a uniform fidelity to the result of the PFG NMR experiments in our dataset. We used the same SAXS-optimized conformational ensembles used in our previous work (produced with the CALVADOS model (57)). In our previous work we used the Kirkwood-Riseman equation(56), two empirical relationships relating *R*_g_ and *R*_h_ (58) and HullRad (59) to calculate *R*_h_ from the conformational ensembles generated by CALVADOS. We here use the more recent HullRadSAS approach (24). When we compare the *R*_h_ values calculated from the SAXS-refined ensembles to the revised experimental values, the results are less clear than when we compared to the *R*_h_ values based on the original reference value for dioxane. Specifically, we now find that the Kirkwood-Riseman systematically underestimates the *R*_h_, whereas the other approaches overestimate *R*_h_ (Fig. 6). Of the four methods examined, we find that HullRadSAS agrees more closely with the data This observation is due to the finding that HullRadSAS giving rise to very good agreement with the experimental values for four of the IDPs; for the remaining the result is more complex as the experimental value lies in between the predictions from the different models (Fig. S6). In light of this result, we suggest the use of HullRadSAS when calculating the *R*_h_ of IDP conformations. Assuming that the SAXS-refined conformational ensembles that we have employed are accurate in reproducing the *R*_h_ of these IDPs, and therefore that the major source of uncertainty comes from the model used to calculate the *R*_h_, we suggest that an uncertainty of 7% is taken into account on the calculated *R*_h_.

**Figure 6:**
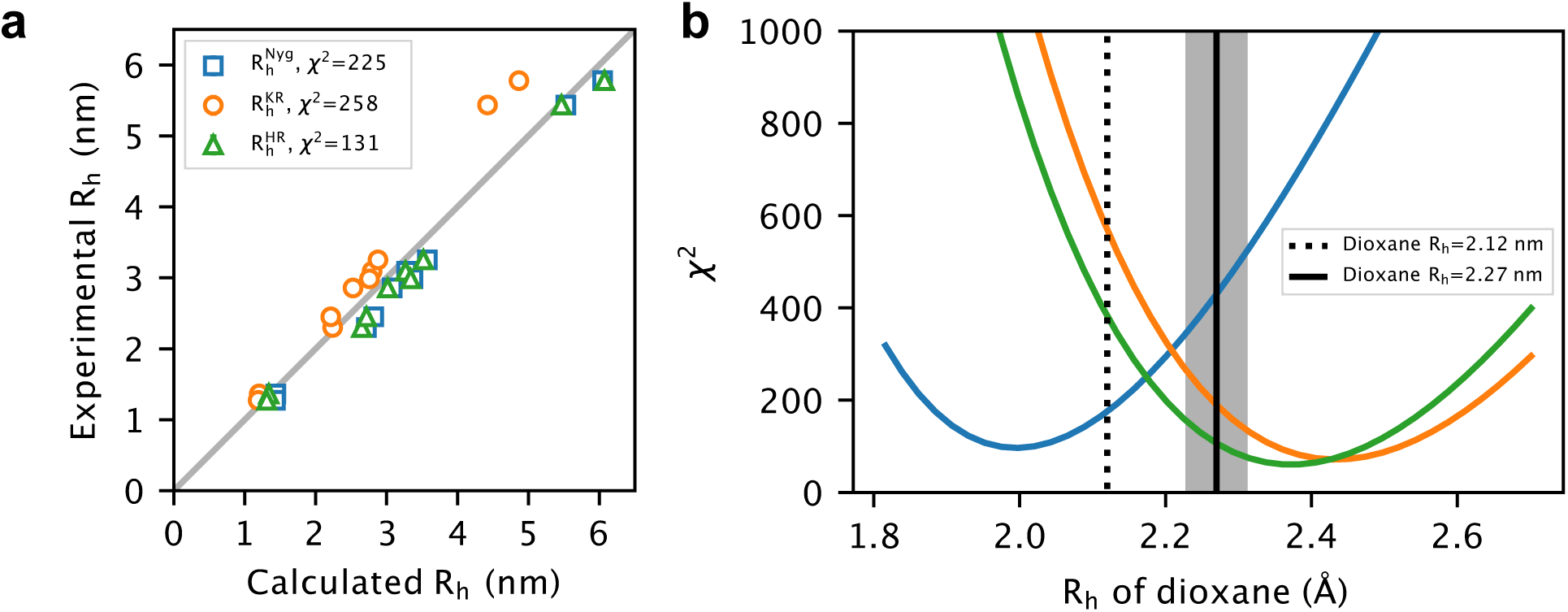
Assessment of models to calculate *R*h from structural ensembles of intrinsically disordered proteins using the same conformational ensembles and approach as in our previous work (10). We compare three approaches to calculate the *R*h from conformational ensembles of several IDPs (Table S4): the Nygaard equation (in blue) (62), the Kirkwood-Riseman equation (in orange) (63) and HullRadSAS (in green) (24). (a) Agreement between calculated and experimentally derived *R*h values. (b) As the value for the *R*h of dioxane varies, so does our assessment of the models to calculate *R*h from conformational ensembles of IDPs: when using a value of 2.12 Å, the Kirkwood-Riseman equation leads to the best agreement with PFG NMR measurements. With the new proposed value of 2.27 ± 0.04 Å, HullRadSAS appears to be the model leading to the best agreement with PFG NMR measurements.

## CONCLUSIONS

From the PFG NMR and SAXS data on the seven proteins, and subsequent weighted average and χ^2^ estimations of the 1,4-dioxane *R*_h_, we find that the original 2.12 Å *R*_h_ is likely underestimated and suggest a value of 2.27 ± 0.04 Å to be used as the standard 1,4-dioxane *R*_h_ when performing PFG NMR to determine the hydrodynamic radius of a protein. The error on this final value is determined as the propagated relative error from technical replications measured by both PFG NMR and SAXS on HEWL and RNaseA; however, this uncertainty will rarely be the limiting factor in the accuracy of derived protein *R*_h_ values. Previously published PFG NMR protein *R*_h_ values can easily be re-referenced to our suggested 1,4-dioxane Rh by increasing the protein *R*_h_ with 7%, however this is only needed as long as absolute *R*_h_ values are reported. While our data suggest that the 1,4-dioxane *R*_h_ is greater than 2.12 Å, one would ideally use an even greater protein dataset and/or other methods to analyse dioxane. Furthermore, the *R*_h_ of 1,4-dioxane might be both pressure– and temperature-dependent, and as such, the *R*_h_ of 1,4-dioxane at different experimental conditions should also be examined further (60, 61).

## Acknowledgements

We thank Signe A. Sjørup for skilled technical assistance and Johan G. Olsen for valuable discussions on water layers in relation to SAXS measurements. We thank Andreas Prestel, manager of the cOpenNMR facility https://www1.bio.ku.dk/copennmr/ (grant no. NNF18OC0032996) for NMR assistance. We acknowledge access to the University of Copenhagen small-angle X-ray scattering facility, CPHSAXS, funded by the Novo Nordisk Foundation (grant no. NNF19OC0055857), and thank Pernille Sønderby Tuelung for assistance https://drug.ku.dk/core-facilities/cphsaxs/. We acknowledge access to computational resources from the Biocomputing Core Facility at the Department of Biology, University of Copenhagen.

## Supplemental figures

**Supplemental figure 1:**
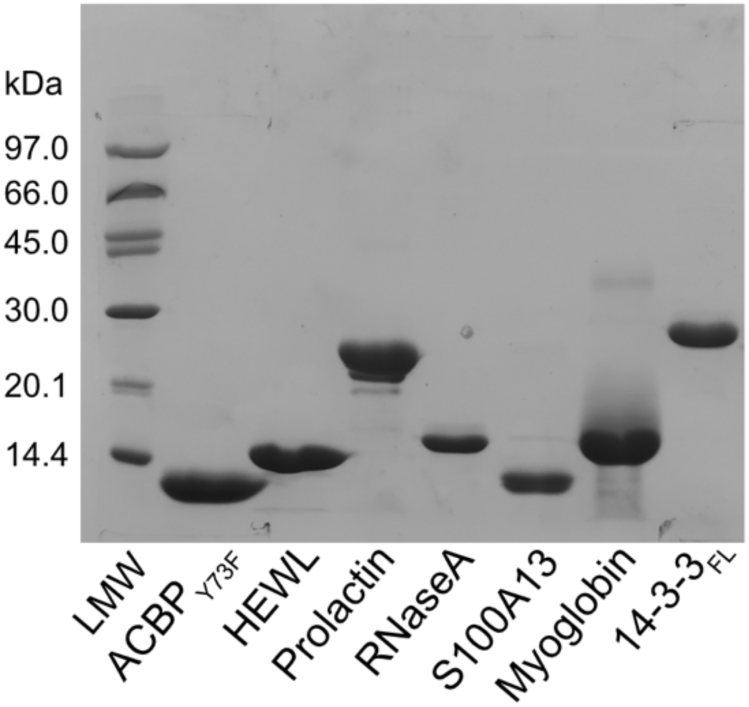
SDS-PAGE of the seven model proteins. The gel has been overloaded with protein to increase visibility of any impurities.

**Supplemental figure 2:**
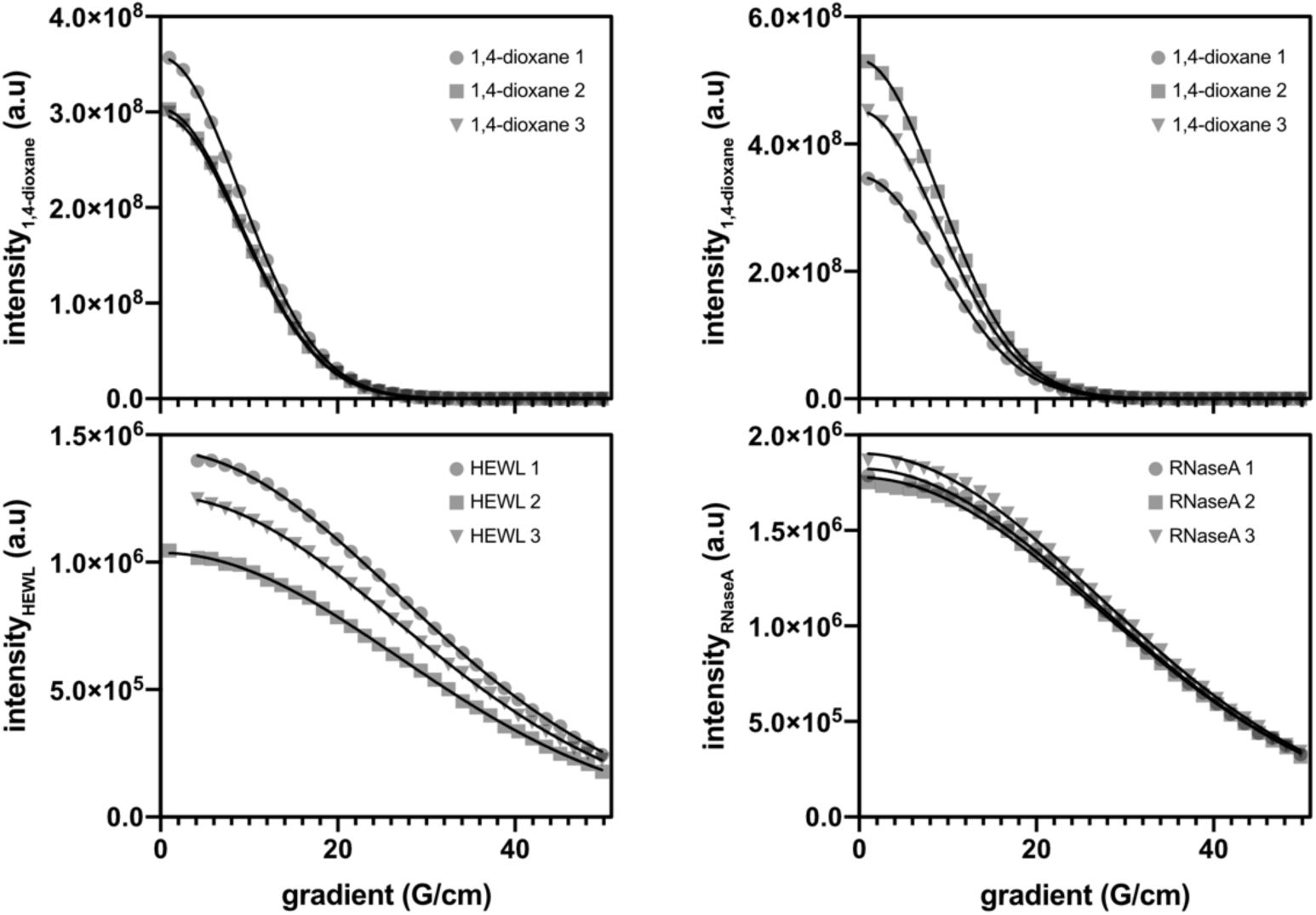
NMR peak intensity decays as a function of gradient strength. Intensity decays are shown for triplicate HEWL samples (left) and triplicate RNAseA samples (right) including decays of 1,4-dioxane (top) and proteins (bottom).

**Supplemental table 1:** Diffusion coefficients of triplicate HEWL and RNAseA and their averages, and the corresponding ratio of the diffusion coefficients (1,4-dioxane/protein) for each sample.

|  | HEWL 1 | HEWL 2 | HEWL 3 | HEWL AVERAGE | RNASEA 1 | RNASEA 2 | RNASEA 3 | RNASEA AVERAGE |
| --- | --- | --- | --- | --- | --- | --- | --- | --- |
| **1,4-DIOXANE** |  |  |  |  |  |  |  |  |
| $D_T$<br>($m^2/s$) | $1.09 \times 10^{-9}$<br>$\pm 3.09 \times 10^{-12}$ | $1.08 \times 10^{-9}$<br>$\pm 1.86 \times 10^{-12}$ | $1.08 \times 10^{-9}$<br>$\pm 4.52 \times 10^{-12}$ | $1.08 \times 10^{-9}$<br>$\pm 9.46 \times 10^{-12}$ | $1.06 \times 10^{-9}$<br>$\pm 7.45 \times 10^{-13}$ | $1.09 \times 10^{-9}$<br>$\pm 2.72 \times 10^{-12}$ | $1.08 \times 10^{-9}$<br>$\pm 1.82 \times 10^{-12}$ | $1.08 \times 10^{-9}$<br>$\pm 5.26 \times 10^{-12}$ |
| **PROTEIN** |  |  |  |  |  |  |  |  |
| $D_T$<br>($m^2/s$) | $1.22 \times 10^{-10}$<br>$\pm 3.59 \times 10^{-13}$ | $1.21 \times 10^{-10}$<br>$\pm 4.71 \times 10^{-13}$ | $1.22 \times 10^{-10}$<br>$\pm 2.74 \times 10^{-13}$ | $1.22 \times 10^{-10}$<br>$\pm 1.10 \times 10^{-12}$ | $1.20 \times 10^{-10}$<br>$\pm 6.24 \times 10^{-13}$ | $1.20 \times 10^{-10}$<br>$\pm 6.18 \times 10^{-13}$ | $1.19 \times 10^{-10}$<br>$\pm 6.29 \times 10^{-13}$ | $1.20 \times 10^{-10}$<br>$\pm 1.87 \times 10^{-12}$ |
| **DIFFUSION RATIO**<br>(relative error) | $8.96 \pm 0.05$ | $8.88 \pm 0.05$ | $8.84 \pm 0.06$ | $8.89 \pm 0.11$<br>(1.3%) | $8.87 \pm 0.05$ | $9.03 \pm 0.07$ | $9.07 \pm 0.06$ | $8.99 \pm 0.15$<br>(1.6%) |

**Supplemental table 2:** Guinier derived *R*g with standard error of the mean from triplicate SAXS measurements of HEWL (2mg/ml) and RNAseA (3mg/ml).

|  | HEWL 1 | HEWL 2 | HEWL 3 | HEWL AVERAGE | RNASEA 1 | RNASEA 2 | RNASEA 3 | RNASEA AVERAGE |
| --- | --- | --- | --- | --- | --- | --- | --- | --- |
| **PROTEIN** |  |  |  |  |  |  |  |  |
| $R_g$ (Å)<br>(relative error) | $14.85 \pm 0.19$ | $15.25 \pm 0.17$ | $15.14 \pm 0.17$ | $15.08 \pm 0.18$<br>(1.2%) | $15.44 \pm 0.15$ | $15.35 \pm 0.15$ | $15.34 \pm 0.16$ | $15.38 \pm 0.15$<br>(1.0%) |

**Supplemental figure 3:**
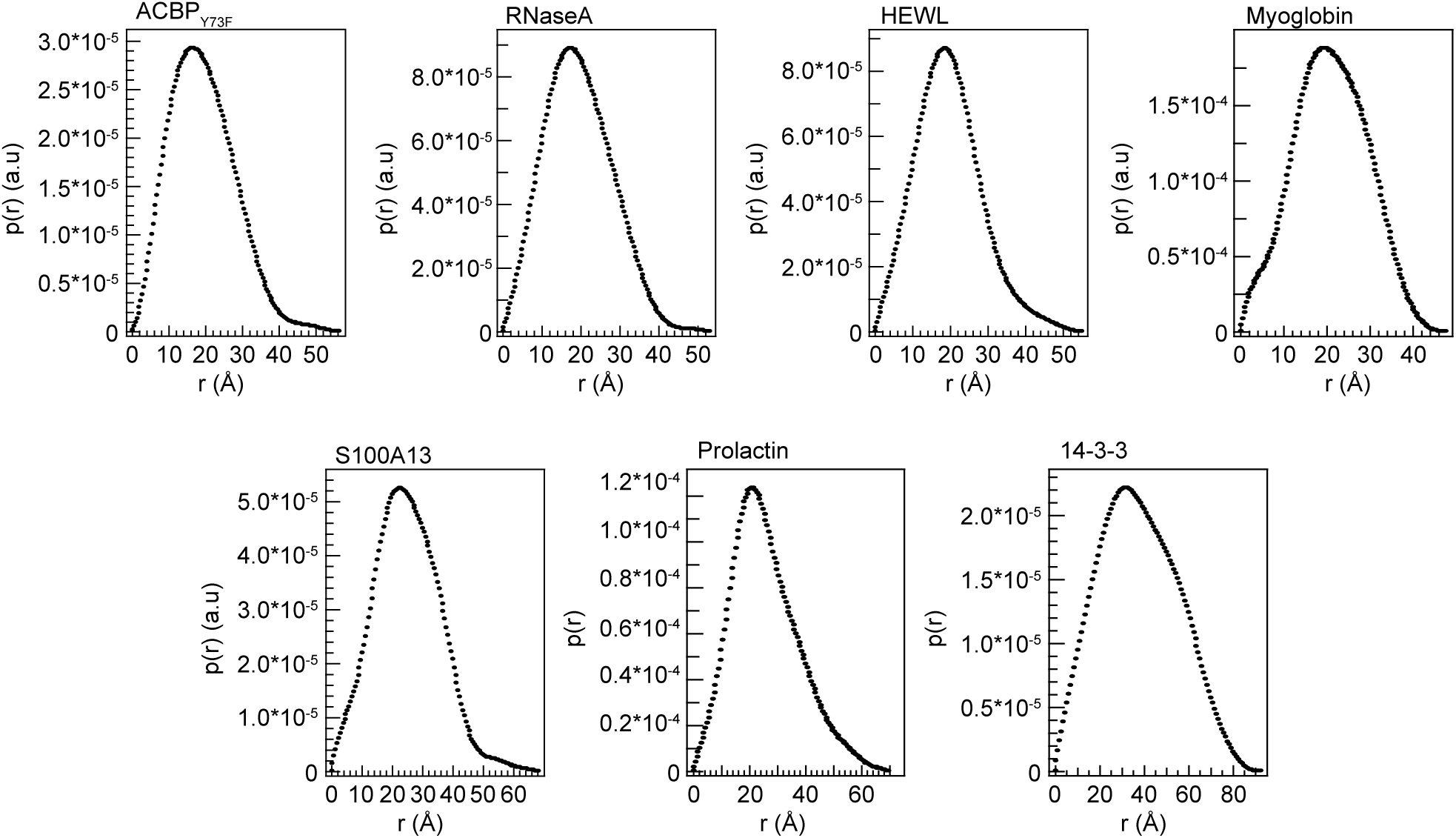
Pair-distance distribution plots (p(r)) of the consensus scattering curve for each of the seven proteins.

**Supplemental table 3:**
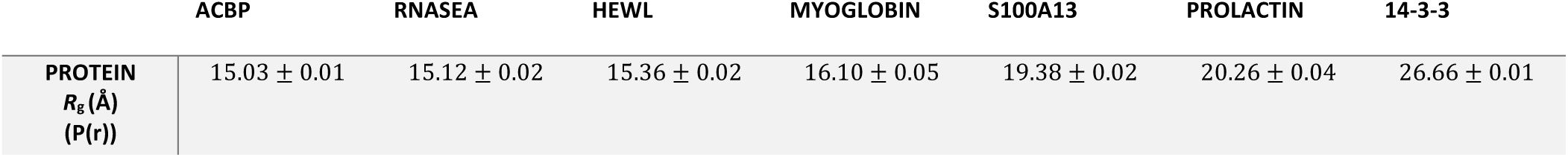
Experimentally derived *Rg* values by pair-distance distribution plots from batch SAXS. All values are reported with propagated standard errors of fit.

**Supplemental figure 4:**
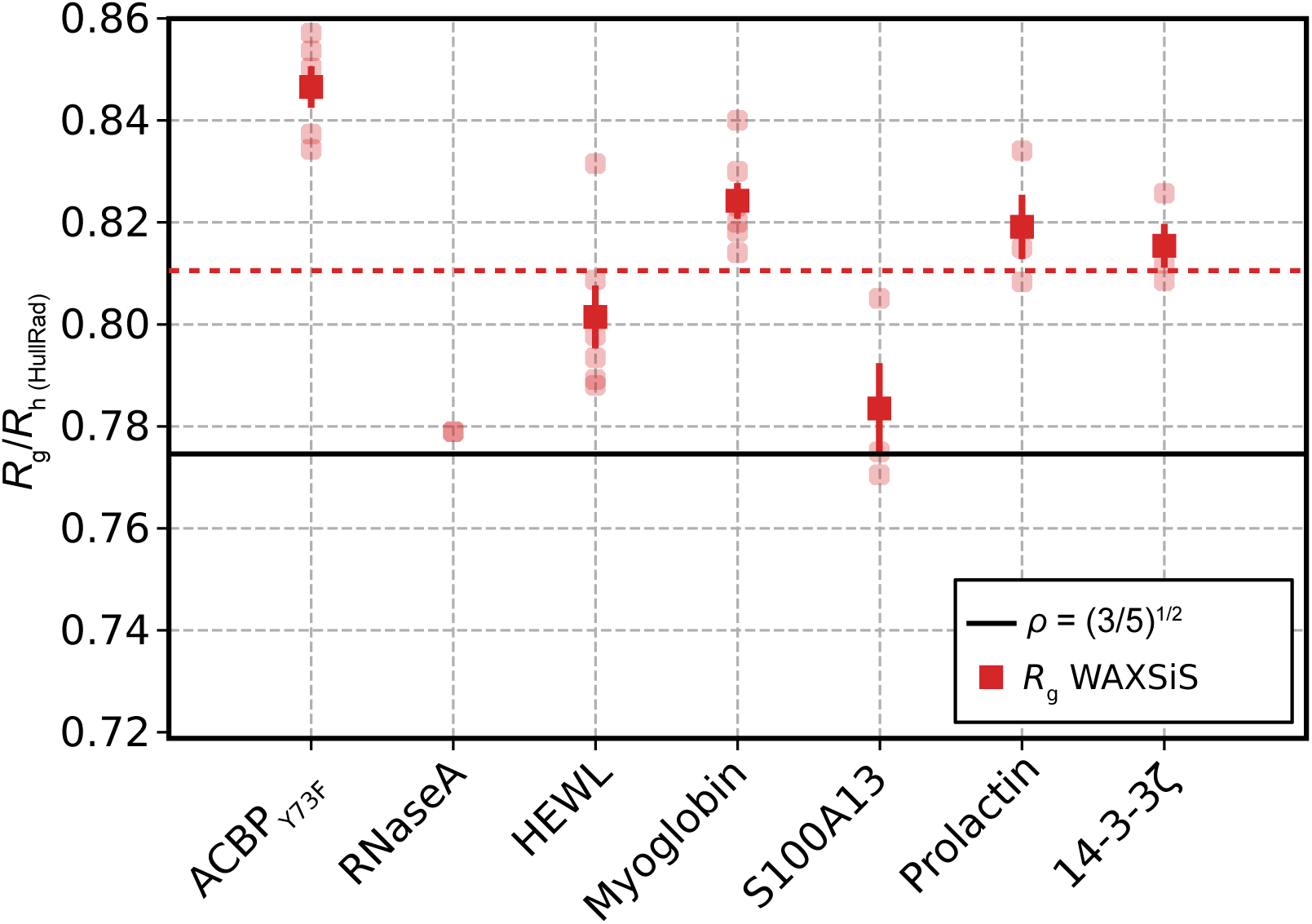
Ratios of *R*g/*R*h obtained from 3D structures using an anhydrous Rg calculation and adding a water shell. For each protein, three to five 3D structures were used, with the following PDB codes for each protein: HEWL (1e8l (28), 1dpx (34), 6abn (35), 5a3e (36)), RNaseA (2aas (27), 1fs3 (37), 1jvt (38), 4ooh(39), 1kf7 (40)), myoglobin (5zze (29), 1wla (41), 4dc8 (42), 5d5r (43), 5cn4 (43)), S100A13 (1yuu (30), 2h2k (44), 2egd (45)), ACBP (1nti (20), 1hb6 (46), 1hb8 (46), 2abd (47)), prolactin (1rw5 (31), 2q98 (48)), 14-3-3ζ (2o02 (32), 1qja (49), 1gjb (50)). AlphaFold structures (51, 52) were also included for each protein. Individual ratios are shown as partially transparent markers. For each protein we calculated the mean ration over different structures and the standard error of the mean (solid markers). Dashed red line represents the calculated average value of the *R*g/*R*h.

**Supplemental figure 5:**
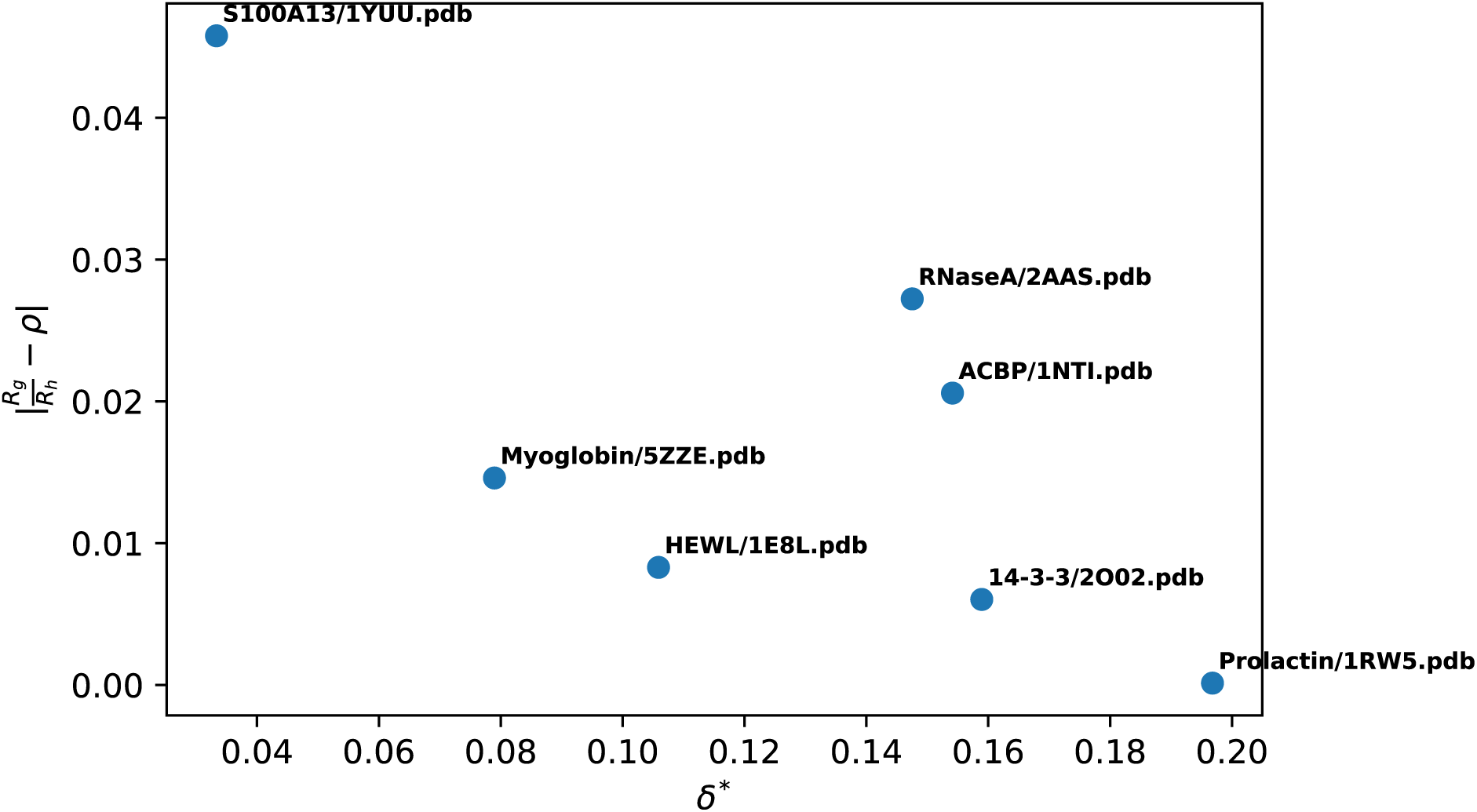
Calculation of the asphericity (d*, as defined in (33) for selected structures of the seven proteins and plot it against the absolute value of the difference between their Rg/Rh ratio to π.

Supplemental table 4: PFG NMR derived *R*h corrected according to a 1,4-dioxane *R*h of 2.27 ± 0.04 Å.

| IDP | Hst5 | RS | DSS1 | Sic1 | ProTa | NHE6-cmdd | A1 | aSyn | ANAC-046 | GHR-ICD | Tau |
| --- | --- | --- | --- | --- | --- | --- | --- | --- | --- | --- | --- |
| $R_h$ (Å) corrected | 1.36 | 1.27 | 1.82 | 2.3 | 3.1 | 2.86 | 2.45 | 2.985 | 3.25 | 5.4 | 5.8 |
| $R_h$ error (Å) corrected | 0.04 | 0.02 | 0.07 | 0.1 | 0.1 | 0.05 | 0.08 | 0.006 | 0.06 | 0.1 | 0.2 |

**Supplemental figure 6:**
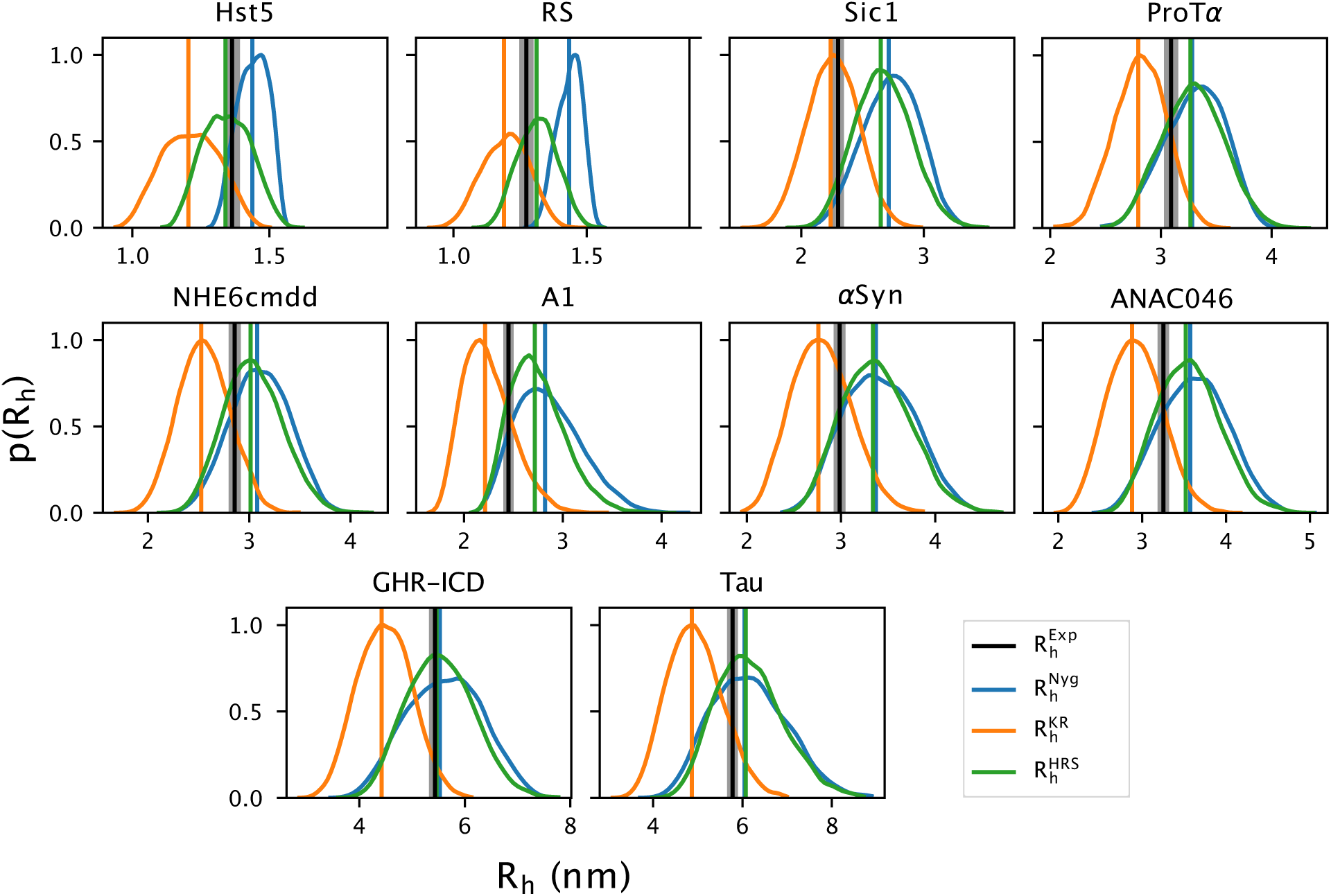
Probability distributions of the *R*h and their ensemble averages calculated from SAXS-reweighted CALVADOS ensembles, compared with the *R*h determined by PFG NMR diffusion (in black). We tested four approaches to calculate the *R*h from atomic coordinates: the *R*g-dependent Nygaard equation (Nyg, in blue), the Kirkwood-Riseman equation (KR, in orange), HullRadSAS (HRS, in green).

